# Ecosystem dynamics in wet heathlands: spatial and temporal effects of environmental drivers on the vegetation

**DOI:** 10.1101/2024.08.06.606794

**Authors:** Christian Damgaard

**Author notes:** Corresponding author: Christian Damgaard.

## Abstract

To understand and estimate the effects of environmental drivers on wet heathland vegetation, pin-point cover data from 42 sites sampled during a 15-year period was regressed onto environmental variables (nitrogen deposition, soil pH, soil C-N ratio, soil type, precipitation and grazing) in a spatio-temporal structural equation model using a Bayesian hierarchical model structure with latent variables to model the effect of measurement and sampling uncertainties.

The results suggest that the modelled environmental variables have various regulating effects on the large-scale spatial variation as well as plant community dynamics in wet heathlands. Most noticeably, nitrogen deposition and yearly precipitation had relatively large and opposite effects on the characteristic species *Erica tetralix* and *Molinia caerulea*, where nitrogen deposition had negative effects on *E. tetralix* and positive effects on *M. caerulea*. The results of this study differed in important qualitative aspects from the findings of an earlier study where comparable data from a shorter time-series (7 years instead of 15 years) were analyzed with a similar model, which suggests that relatively long time-series are needed for studying ecosystem dynamics. Furthermore, it was concluded that the effect of nitrogen deposition on plant community dynamics mainly was through direct effects, whereas the effect of soil type on plant community dynamics was both direct and indirect mediated by the effect of soil type on soil pH.

It was concluded that the modeled environmental variables are sufficient for predicting the *average* plant community dynamics of wet heathlands. However, caution are required if the fitted model is used for generating local ecological predictions as input to a process of generating adaptive management plans for specific wet heathland sites. Moreover, the results suggest that the ratio between the two species *E. tetralix* and *M. caerulea* may be used as an indicator for the conservation status of wet heathlands.

## Introduction

To appropriately manage semi-natural wet heathland ecosystems under the influence of climate change and altered land use practices, it is important to understand and be able to predict the effects of environmental drivers on the vegetation. Overall, the conservation status of Danish wet heaths is deteriorating and is with high certainty unfavorable in more than 40% of the areas (Nygaard et al. 2020). Improved quantitative understanding of the effects of the different environmental drivers will enable us to make quantitative predictions at the local site level to future ecosystem responses to changes in e.g. climate and grazing regimes as well as provide important input for local adaptive management plans (Damgaard 2021). Consequently, to better understand the causal relationships underlying the observed large-scale variation and temporal changes in wet heathlands as well as estimate the effect of the environmental drivers, a structural equation model of the wet heathland ecosystem was hypothesized and fitted to spatial and temporal ecological data from 42 Danish wet heaths in the period 2007 to 2021. Generally, the effects of changes in environmental variables, e.g. climate, soil physical properties and disturbance regimes, on plant communities are expected to occur with some time-lags (e.g. Svenning and Sandel 2013). Furthermore, the regulating environmental variables are expected to vary considerably among different sites, and it is critical to integrate this large-scale spatial variation into the analysis of the ecosystem dynamics.

The *a priori* selection of the studied environmental drivers was based on current ecological knowledge, where specific environmental drivers, e.g. atmospheric nitrogen deposition, soil type, soil pH and disturbance, have been shown to have an effect on wet heaths vegetation (e.g. Aerts and Berendse 1988; Damgaard et al. 2013; Grant et al. 1996; Lykke et al. 2015). However, wet heathland vegetation is expected to be regulated by more environmental drivers than it was possible to include in this study because of missing data, e.g. hydrology, species-specific herbivores and pests and previous natural management actions. Some of these more or less unknown factors can have a geographical regional structure that may be partly explored using latent geographic factors (Ovaskainen et al. 2016), and possible significant effects of such latent geographic factors can generate new testable causal hypotheses. Furthermore, it is expected that soil type and nitrogen deposition may have both direct and indirect effects by affecting soil pH (Damgaard et al. 2014), and such direct or indirect causal pathways are often best modelled using structural equation models (SEM) that are fitted to observed ecological data (Grace et al. 2010). Generally, a SEM does not allow us to prove the hypothesized causal relationships, but it is possible to test whether specific casual relationships are supported by data. To demonstrate causality, it is necessary to manipulate the system, e.g. in the form of a manipulated experiment, and observe whether the response predicted by the SEM actually takes place (Granger 1969; Pearl 2009).

The dynamics of semi-natural heathland ecosystems, and especially *Calluna* dominated heaths, were first studied by Watt (1947), who gave a detailed account of possible ecological processes that lead to different spatial vegetation patterns. This line of work has since been extended by several authors (e.g. Gimingham 1978; Gimingham 1988; Gimingham et al. 1981; Løvschal and Damgaard 2022; Usher and Thompson 1993), who describe the spatial patterns of the heathland vegetation at different scales and how they are regulated by management. More specifically, the large-scale spatial distribution of *Erica tetralix* in Danish wet heaths is negatively correlated with nitrogen deposition and acidification (Damgaard et al. 2014), and *E. tetralix* displayed positive large-scale covariance with *Calluna vulgaris*, which again was negatively correlated with *Molinia caerulea* (Damgaard et al. 2017). Furthermore, a significant decline in the cover of *E. tetralix* at Danish wet heaths has been observed (Damgaard 2012).

The objective of the present study is to update the findings in a previous study of Danish wet heathland ecosystems (Damgaard 2019). In the first study, ecological monitoring data from the period 2007 - 2014 were analyzed, but now more data have been collected, and it is interesting to investigate to which extend the previous findings may be confirmed and whether new conclusions may be extracted from the analysis of the more comprehensive data set. In the previous study, the multi-variate relative cover of four species groups was modelled (*Erica tetralix, Calluna vulgaris, Molinia caerulea* and an aggregated class of all other plants), and this list has now been extended to six species groups by dividing the class of all other plants into all other graminoids, all other herbs and cryptogams. Furthermore, the modelled list of environmental variables in the first study (nitrogen deposition, soil type, soil pH, precipitation and grazing) has now been extended with the C-N ratio in the topsoil. The main conclusions of the first study (Damgaard 2019) were that the two dwarf shrubs, *E. tetralix* and *C. vulgaris*, responded in qualitatively similar ways to the abiotic variables and qualitatively oppositely to the grass *M. caerulea* and the aggregate class of other plants. More specifically, the two dwarf shrub species most likely were positively affected by nitrogen deposition, soil pH, sandy soils, low precipitation and the absence of grazing.

## Materials and Methods

### Wet heathlands

Wet heathlands are semi-natural ecosystems on sandy, nutrient-poor soils with a natural high-water table, which first became widespread in prehistoric times under the influence of extensive agricultural practices (Løvschal 2021). In Denmark, the wet heathlands are mainly situated on the relatively flat sandy nutrient-poor soils in the western part of Denmark that were not covered by ice during the last (Weichselian) glacial period (Fig. 1). The vegetation is mainly comprised of dwarf shrubs and graminoids (Hampton 2008), where especially *Erica tetralix, Calluna vulgaris* and *Molinia caerulea* are characteristic and often dominating, with *E. tetralix* and *M. caerulea* being more abundant in more humid soils (Gimingham et al. 1979; Rutter 1955; Stace 1999; Usher and Thompson 1993).

**Fig. 1.**
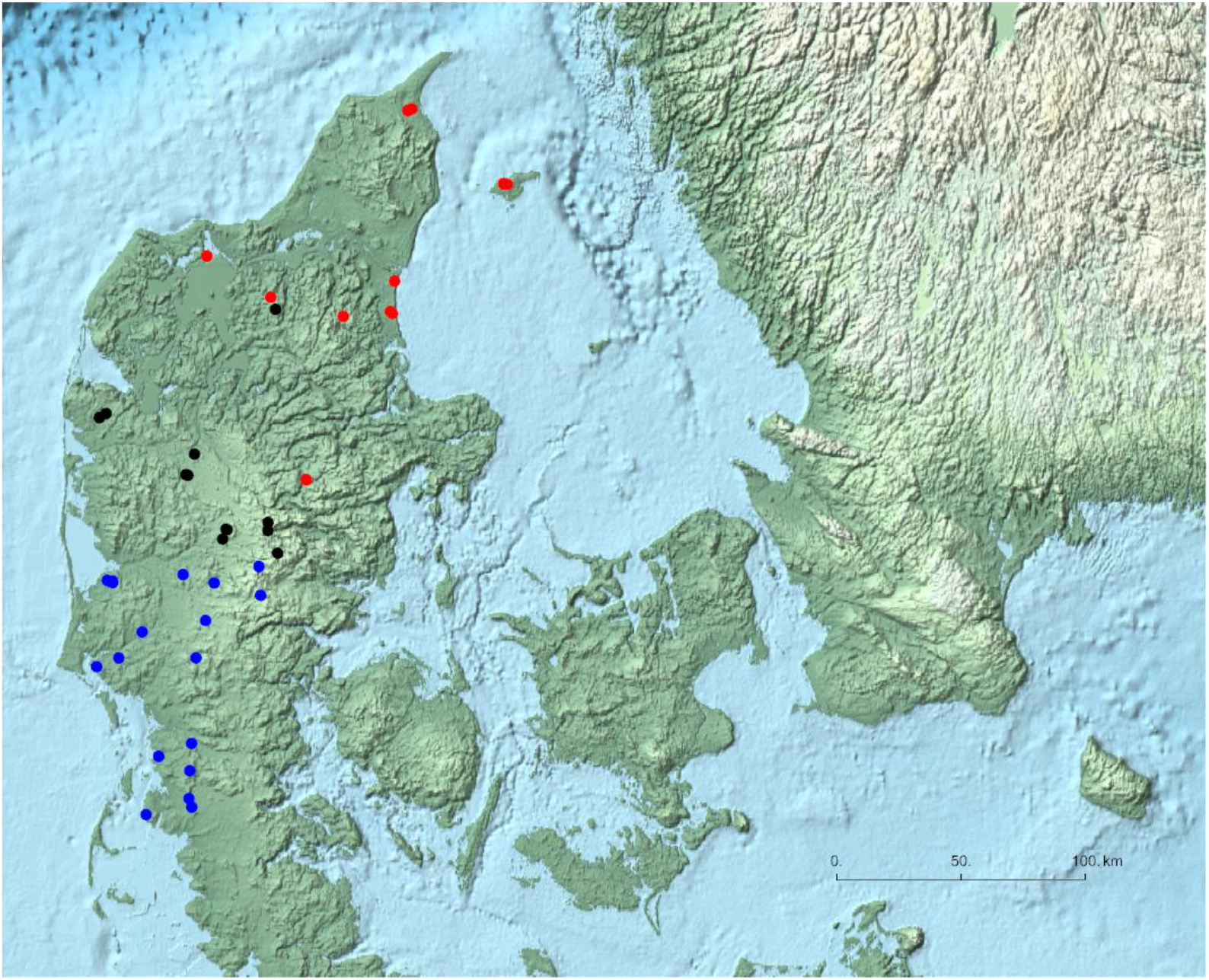
Map of the selected 42 Danish wet heathland sites. Most sites are situated on the penisula Jutland, and the remaining sites are situated on islands in Kattegat. The different colors represent a classification of the different sites into three geographical regions.

*Erica tetralix* (L.) (cross-leaved heath) grows to a height of 10 – 30 cm on acid humid substrate. Plant growth is erect and straggling when competing with other species (Bannister 1966). In wet habitats, it has an almost creeping lifeform and the lower branches are often buried in moss or litter and can produce roots and shoots, whereas in drier soil the species tends to grow in tussocks (Bannister 1966).

*Calluna vulgaris* (L.) Hull (heather) is a 10 to 50 cm tall dwarf shrub characteristic of sandy and peaty heathland soil. *Calluna* is often dominant on heaths, moors and nutrient-poor grasslands, though not tolerant to heavy grazing, and prefers at least moderately well drained soils (Gimingham 1960). The developmental phases of heather include pioneer, building, mature and degenerate (Usher and Thompson 1993; Watt 1947), and without management actions such as burning, grazing or cutting, *Calluna* plants degenerate when their age exceeds approximately 30 years, thereby leaving room for heathland succession.

*Molinia caerulea* (L.) Moench (purple moor grass) is a deciduous, tussock-forming grass with swollen basal internodes and roots acting as overwintering storage organs (Grant et al. 1963). The grass preferably grows at sites with ground-water movement, good soil aeration and enriched nutrient supply as a calcifuge and typically grows to a height of 95 cm, culm and panicle included (Stace 1999; Taylor et al. 2001). The species is resistant to burning (Grant et al. 1963; Grant et al. 1996).

The estimated cover and change in cover of the most common species in Danish wet heathland plots are shown in Table S1.

### Sampling design

Hierarchical time-series data from 42 wet heathland sites (Fig. 1) that had been monitored at least three times in the period from 2007 to 2021 were used in the analysis. Twenty-three of the 42 sites are NATURA 2000 habitat sites and are protected under the habitat directive (EU 1992). All sites included several plots classified as wet heathland (EU habitat type: 4010) according to the habitat classification system used for the European Habitat Directive EU (EU 2013). The area of the sites ranged from 0.5 ha to 133 ha, with a median area of 9.5 ha. A total of 353 unique plots were used in the analysis. The sampling intensity was irregular among sites and years, but typically ten plots were sampled from each site at each time point. Including resampling over the years, a total of 1450 plot data were used in the analyses. The plots were resampled with GPS-certainty (< 10 meters).

The data are a subset of the data collected in the Danish habitat surveillance program NOVANA (Nielsen et al. 2012; Nygaard et al. 2024).

### Variables and measurements

#### Plant cover data

The plant cover, which is the relative projected area covered by a species, was measured for all higher plants by the pin-point method using a square frame (50 cm X 50 cm) of 16 grid points that were equally spaced by 10 cm (Nielsen et al. 2012). At each grid point, a thin pin was inserted into the vegetation and the plant species that were touched by the pin were recorded and used as an estimate of cover (Damgaard and Irvine 2019; Levy and Madden 1933; Lindquist 1931). Since the pin-point cover data after 2007 were recorded for each pin separately, the species cover data are readily aggregated into cover data for classes of species at a higher taxonomic or functional level. At each grid point, the pin may hit different plant species from the same species class and, in those cases, the hits are only counted as a single hit of the species class at the grid point.

In this study, the species were classified into six species groups: *Erica, Calluna, Molinia*, any other graminoids, any other herbs or a cryptogam. The assumed distribution of pin-point cover data for single species and the joint distribution for multiple species are outlined in the electronic supplement (Appendix A).

#### Nitrogen deposition

Nitrogen deposition at each plot was calculated for each year using a spatial atmospheric deposition model in the period from 2005 to 2014 (Ellermann et al. 2012). The mean site nitrogen deposition ranged from 7.34 kg N ha^−1^ year to 23.75 kg N ha^−1^ year^−1^, with a mean deposition of 14.09 kg N ha^−1^ year^−1^. Anthropogenic nitrogen deposition has reached a maximum and is currently decreasing in Denmark (Ellermann et al. 2018).

#### Soil pH

Soil pH was measured in randomly selected plots from the uppermost 5 cm of the soil (four samples were amassed into a single sample). The soils were passed through a 2mm sieve to remove gravel and coarse plant material, and pH_KCl_ was measured on a 1 M KCl-soil paste (1:1). The soil sampling intensity was irregular among sites and years, but typically between one and four plots were sampled from each site at each time point. When a plot was resampled, the pH at the plot was calculated as the mean of the samples. In total, 632 independent soil pH values were used in the analysis. The measured soil pH ranged from 2.7 to 6.7, with a mean soil pH of 3.33.

#### C-N ratio in the soil

Soil C-N ratio was measured in randomly selected plots from the uppermost 5 cm of the soil (four samples were amassed into a single sample). Total C in each sample was determined by dry combustion and N by the Kjeldahl method. The soil sampling intensity was irregular among sites and years, but typically between one and four plots were sampled from each site at each time point. When a plot was resampled, the C-N ratio at the plot was calculated as the mean of the samples. The measured C-N ratio at the site level ranged from 16.97 to 36.33, with a mean site C-N ratio of 25.89. Moreover, the average soil C–N ratio on wet heathlands has been observed to decrease in the period from 2004 to 2014 (Strandberg et al. 2018).

Since soil C-N ratio is expected to be influenced by the vegetation, e.g. by the amount of slowly decomposing dwarf shrub litter, soil C-N ratio may be thought of more as an environmental covariable than an environmental driver.

#### Soil type

The texture of the topsoil for each site was obtained from a raster based map of Danish soils (Greve et al. 2007). The categorical classification of the soil (JB-nr.) was made on an ordinal scale with decreasing particle size, 1: coarse sandy soil, 2: fine sand soil, 3: coarse loamy soil, 4: fine loamy soil. There were some records with other soil types, but because of possible classification errors they were treated as missing values. The mean soil type was 1.35.

#### Precipitation

Site-specific precipitation was measured by the average annual precipitation in the period 2001 to 2010, with a spatial resolution of 10 km (DMI 2014). The annual precipitation ranged from 653 mm to 962 mm, with a mean precipitation of 837 mm.

#### Grazing

Land-use was summarized by possible signs of grazing, e.g. the presence of livestock or short vegetation within fences was recorded by the observer at each plot for each sampling year since 2007 as a binary variable (sign of grazing = 1, no sign of grazing = 0), i.e. if grazing was 0.5, then this probability may arise by a number of ways, e.g. if half the plots at the site showed signs of being grazed each year or all plots were grazed every second year. The mean grazing variable ranged from 0 to 0.93 among sites, but most sites had no grazing and the mean grazing intensity at the site level was 0.15. Unfortunately, the grazing variable does not include information on which animals were used for grazing, stocking densities or grazing duration, and is therefore a quite imprecise variable that must be interpreted together with general knowledge on the typically used grazing regime of wet heathlands.

#### Geographic regions

The 42 wet heathland sites were grouped into three geographic regions (Fig. 1) that were used to investigate possible latent geographic factors.

### Spatio-temporal modelling

It was decided to fit the spatio-temporal structural equation model (SEM) within a Bayesian hierarchical model structure using latent variables to model the effect of measurement and sampling uncertainties (Fig. 2). This use of a hierarchical model structure is important, since it has been demonstrated that ignoring measurement and sampling uncertainties may lead to model and prediction bias (Damgaard 2020; Damgaard and Weiner 2021). Furthermore, it is an advantage when making ecological predictions to separate measurement and sampling uncertainties from process uncertainty. The hierarchical SEM approach and the motivation for using it are explained further in Damgaard (2019), where a similar model was fitted to ecological monitoring data, and more generally in Damgaard (2022). The mathematical and statistical details of the spatio-temporal modelling are explained in the electronic supplement (Appendix B)

**Fig. 2.**
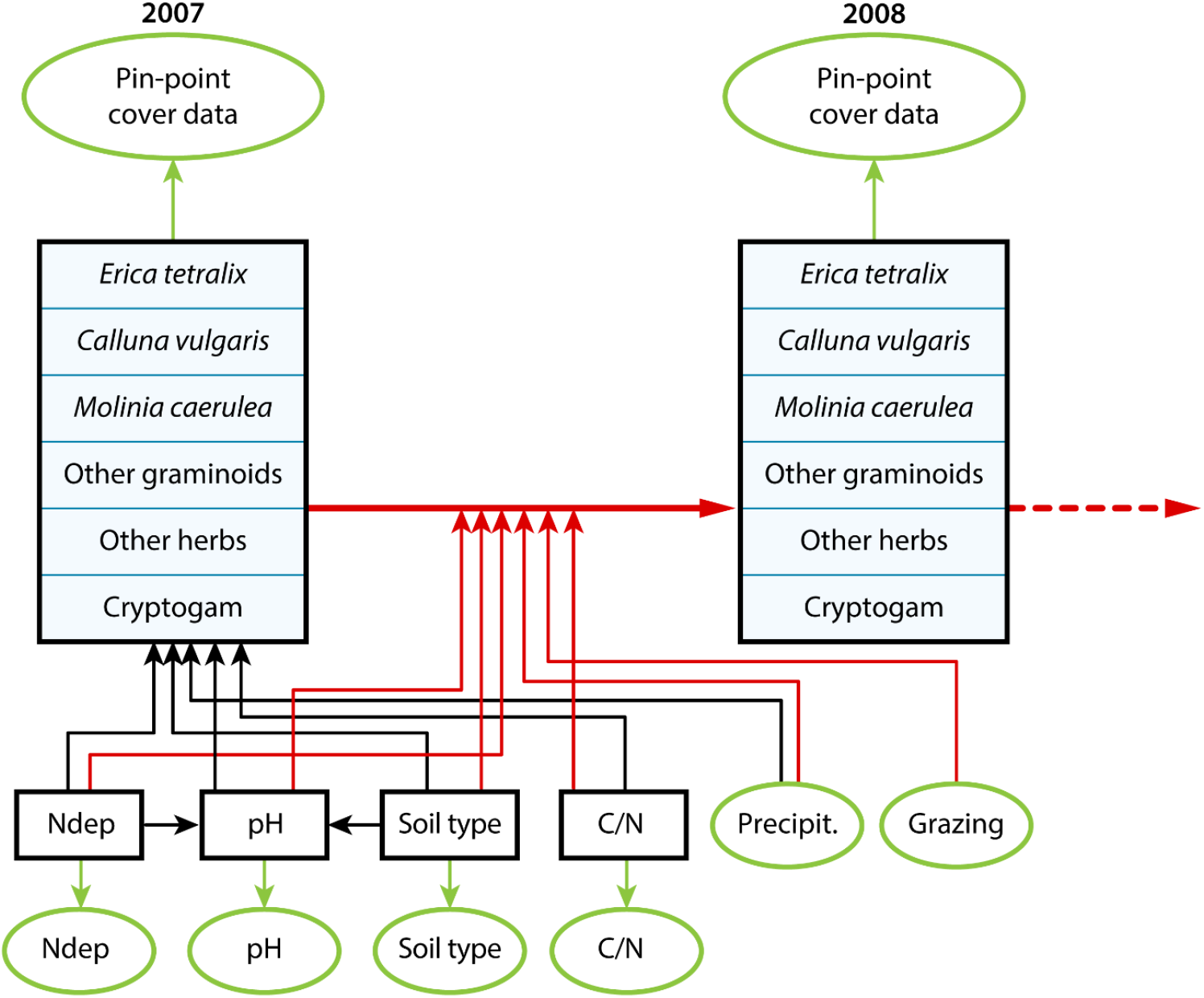
Outline of hierarchical SEM. The spatial variation in vegetation cover in 2007 is modelled by nitrogen deposition (Ndep), soil pH (pH), soil C-N ratio (C/N), soil type and precipitation (Precipit.). The yearly change in vegetation cover from 2007 to 2021 (only a single yearly change is shown in the figure) is modelled by all the former variables as well as grazing. The black boxes are latent variables and the green ovals are data. The black arrows denote large-scale spatial processes, and the red arrows denote temporal processes (Appendix B).

The procedures for estimating the most important single species cover and change in species cover for all sampled wet heathland plots since the beginning of the monitoring program in 2004 are explained in the electronic supplement (Appendix C), and the used code and additional tests of fitting properties etc. may be found in Appendix D.

## Results

The estimated cover of the most common species in Danish wet heathlands and the estimated change in cover are shown in Table S1. Notably, the six most abundant species all showed significant changes in cover from the period 2004 to 2021. The cover of the most abundant species, the grass *M. caerulea*, increased significantly, whereas the cover of the five next abundant species (the dwarf shrubs: *C. vulgaris, Empetrum nigrum*, and *E. tetralix*, and the graminoids *Carex nigra* and *Deschampsia flexuosa*) all decreased significantly.

To further understand these observed changes in species composition, vegetation changes were fitted to site variation in selected abiotic and land-use environmental variables in a spatio-temporal structural equation model (Fig. 2). The selected environmental variables covaried at the 42 wetland heath sites (Fig. S1), which again is expected to lead to covariance among parameter estimates and affect the fitting properties of the model negatively. Nevertheless, plots of the mean latent vs. expected logit-transformed cover variables demonstrated a relatively good fit of the large-scale spatial variation in cover (Fig. 3A; between 44% and 68% of the variation is explained, Table S3), and the model fitted the temporal process of the change in cover very well (Fig. 3B; between 88% and 99% of the variation is explained, Table S3). Furthermore, the Dunn–Smyth residuals of the marginal observed cover data of the six species classes were approximately normally distributed (Fig. S2).

**Fig. 3.**
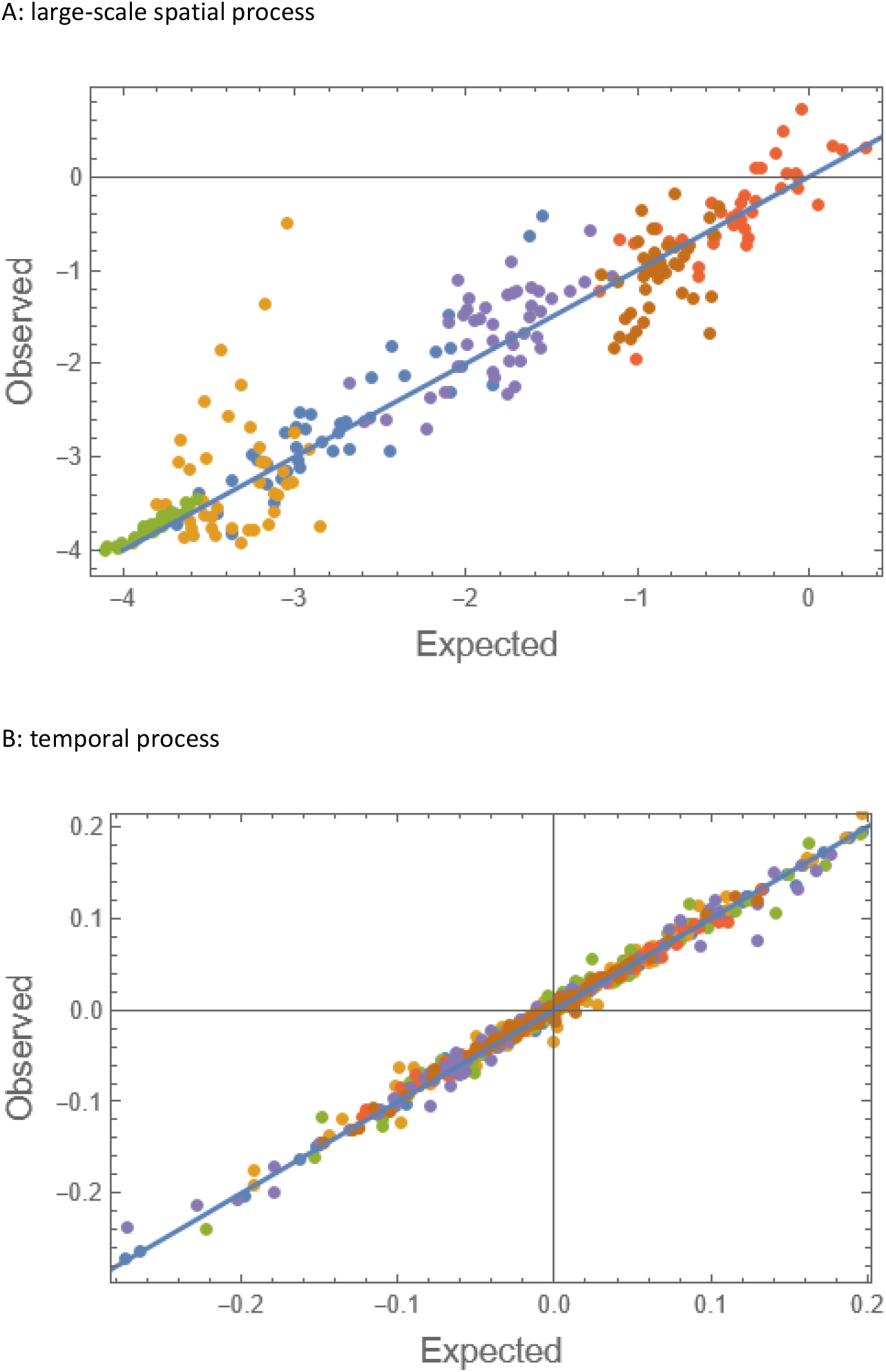
Plots of observed vs. expected logit-transformed cover of the large-scale spatial process (A) and the temporal (B). Blue: *Erica*, yellow: *Calluna*, green: *Molinia*, red: other higher plants.

To prevent possible prediction bias, the different sources of uncertainty, i.e. measurement and sampling uncertainty when measuring plant cover, nitrogen deposition, soil pH, soil C-N ratio and soil type, as well as the structural uncertainties due to the modelled soil pH, large-scale (among sites) spatial variation and the temporal processes, were modelled explicitly. The most important source of measurement uncertainty was the plant cover measurement due to the significant small-scale spatial aggregation of plant species, which was modelled by the parameter δ in the Dirichlet - multinomial mixture distribution. The median estimated value of δ was 0.17 with a relatively narrow credible interval (Table S2). Generally, the estimates of the structural uncertainties were relatively low, although the large-scale spatial variation were relatively high for some species classes (Table S2, Fig. 3A).

Many of the regression parameters that measure the large-scale spatial and temporal effect of the abiotic variables on the vegetation were significantly different from zero (Table S2), suggesting that the modelled environmental and land-use factors have a regulating effect on both the large-scale spatial variation in cover of the six species classes as well as plant community dynamics in wet heathlands. Except for precipitation, all the studied environmental drivers had significant effects on the large-scale spatial variation of two or more species classes (Fig. 4A, Table S2) and, generally, there was significant geographic variation among the three assigned Danish regions (Table S2). Furthermore, except for soil type, all the studied environmental drivers had significant effects on the temporal variation of one or more species classes (Fig. 4B, Table S2). Notably, nitrogen deposition and precipitation had relatively large temporal effects that differed qualitatively among different species classes (Fig. 4B). For example, relatively low nitrogen deposition and high precipitation on average had a positive effect on the cover of *E. tetralix* and other graminoids, whereas the same combination of the environmental drivers had negative effects on the cover of *M. caerulea* and cryptogams (Fig. 4B). Furthermore, this study observed only benign effects of grazing on the plant community dynamics in wet heathlands, except for a significant positive effect on the cover of other herbs (Table S2, Fig. 4B).

**Fig. 4.**
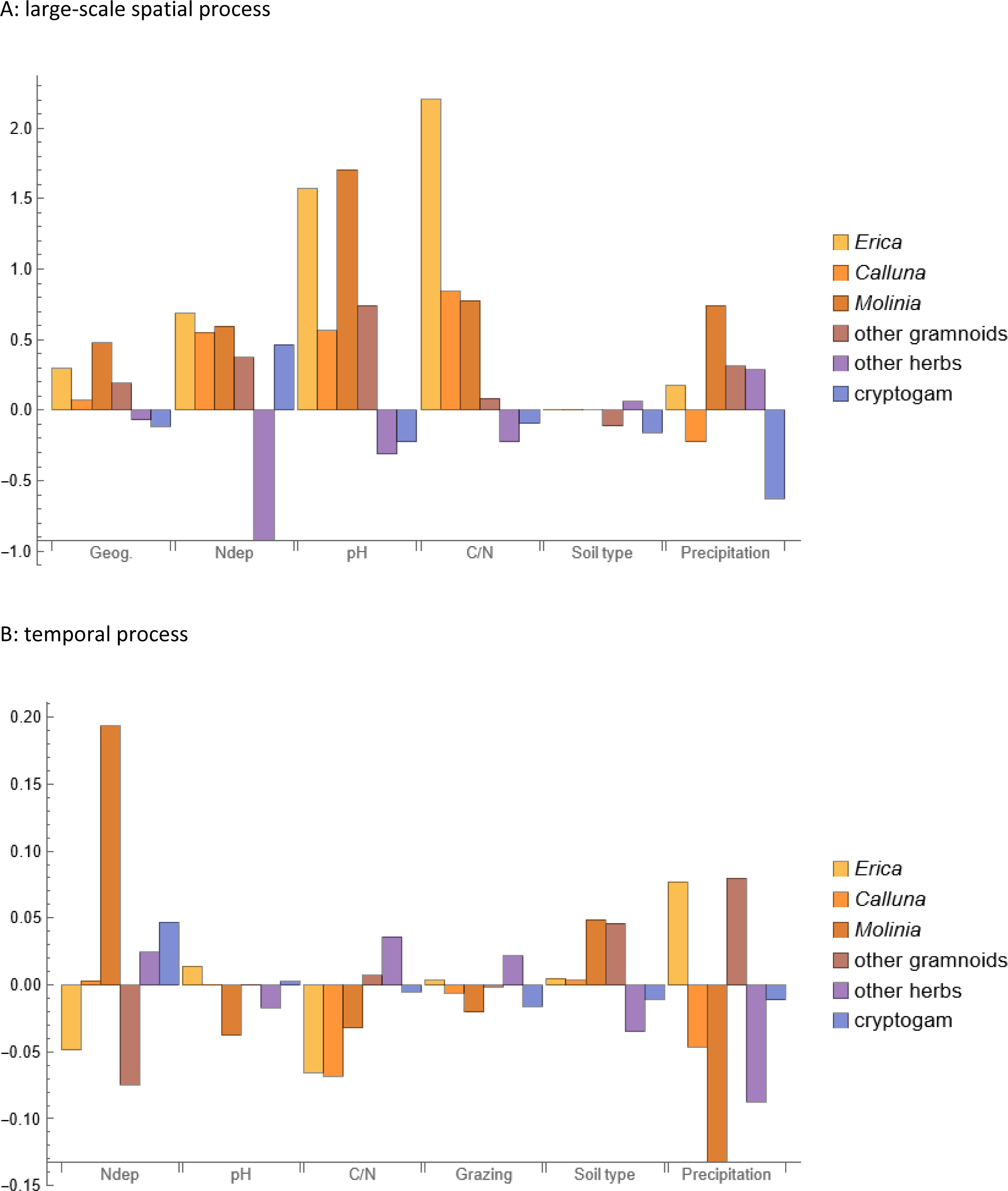
Standardized regression coefficients of the SEM for the large-scale spatial process (A) and the temporal (B).

There were no significant effects of nitrogen deposition on soil pH (*γ*_*N*_, Table S2), but soil pH was found to be significantly lower on more clayey soils compared to sandy soils (*γ*_*S*_, Table S2). It was therefore concluded that the effect of soil type on the vegetation encompassed both direct effects of soil type and indirect effects mediated by the effect of soil type on soil pH. On the other hand, the effect of nitrogen deposition was mainly through direct effects on the vegetation and not by possible soil acidification.

## Discussion

### Environmental drivers and plant community dynamics

Since many of the regression parameters that measure the effect of the environmental drivers on the change in plant species cover were significantly different from zero, the results suggest that the modelled environmental variables have various regulating effects on the large-scale spatial variation as well as plant community dynamics in wet heathlands. The results of this study differed in important qualitative aspects from the findings of the earlier study, where comparable data from a shorter time-series (7 years instead of 15 years) were analyzed with a similar model, which suggests that relatively long time-series are needed for studying ecosystem dynamics.

One or more of the modelled environmental drivers affect the cover of all the six investigated species classes. However, in the following discussion of the results I will focus on the two species *E. tetralix* and *M. caerulea*. Both species are characteristic species of wet heathlands, but in the past decades there has been a dramatic shift in the wet heathland vegetation, such that the tussock forming grass *M. caerulea* has become the dominant species at many sites at the expense of e.g. dwarf shrubs species such as *E. tetralix* (Bobbink et al. 2022; Damgaard 2012). This change has been associated with a considerable deterioration in the biodiversity and conservation status of wet heathland sites. For example, on Randbøl Heath in Southern Denmark, where *M. caerulea* has become dominant in most of the humid parts and has completely displaced *E. tetralix, C. vulgaris* and most other species of the wet heathland (Degn 2006).

The estimated negative effect of nitrogen deposition in the range of 7.34 kg N ha^−1^ year to 23.75 kg N ha^−1^ year^−1^ on the cover of *E. tetralix* corresponds with previous findings in more simple ecological models, where a negative spatial effect on *Erica* abundance has been demonstrated (Damgaard et al. 2017; Damgaard et al. 2014), and a current empirical critical load of wet heathlands between 5 and 15 kg N ha^−1^ year^−1^ (Bobbink et al. 2022). The present results also corroborate the findings of a manipulated experiment in wet heathlands, where the cover of *M. caerulea* increased relative to *E. tetralix* in nitrogen treated plots (Aerts and Berendse 1988), although the manipulated nitrogen level was unrealistically high (200 kg N ha^−1^ year^−1^) compared to current Danish nitrogen deposition levels. Moreover, it was observed that *E. tetralix* was a stronger competitor than *M. caerulea* in control plots, whereas *M. caerulea* was a stronger competitor at unrealistically high nutrient levels of 200 kg N ha^−1^ year^−1^, 40 kg P ha^−1^ year^−1^ and 200 kg K ha^−1^ year^−1^ (Aerts et al. 1990).

Soil acidity is generally considered an important factor in determining the ecological success of plant species (Ellenberg 1979; Pärtel 2002), and in this study it was found that relatively high soil pH was associated with increasing cover of *E. tetralix*, which is in concordance with previous findings using a simpler ecological model (Damgaard et al. 2014). Oppositely, relatively high soil pH was associated with decreasing cover of *M. caerulea*. Furthermore, through the indirect effects of soil texture on soil pH, relatively sandy soils were associated with increasing cover of *E. tetralix* and decreasing cover of *M. caerulea*

The C–N ratio in the topsoil has been suggested as an indicator of the conservation status of heathlands, such that relatively high values indicate a good conservation status (Nygaard et al. 2024). However, in this study we found that a relatively high C–N ratio was associated with decreasing cover of both *E. tetralix* and *C. vulgaris*, which suggests that the C–N ratio may not be an appropriate indicator for the conservation status of wet heathlands. Moreover, it is difficult to sample topsoil under tussocks of *M. caerulea*.

Generally, European wet heathland is found in topographically flat areas with limited drainage and relatively high and constant precipitation (Hampton 2008), and to some extent the average yearly precipitation may be regarded as a proxy for water content in the soil and hydrology. Within wet heathlands, the cover of *E. tetralix* increased significantly at sites with relatively high precipitation, whereas the cover of *M. caerulea* decreased significantly at those sites. These results do not correspond well with the observations that both *E. tetralix* and *M. caerulea* are more abundant on wetter soils (Gimingham et al. 1979; Rutter 1955) and that it is common to observe *C. vulgaris, E. tetralix*, and *M. caerulea*, in that order, on small topographic slopes (Strandberg et al. 2012), indicating that *M. caerulea* may prefer a relatively more humid environment. The annual precipitation in the future climate in Denmark is predicted to increase, but with decreasing summer precipitation and longer summer drought periods (DMI 2017). Based on the present study, it is uncertain how this combination of more extreme weather will influence the future wet heathland vegetation.

There were no significant effects of grazing on *E. tetralix* or any of the other species classes, except for any other herbs where grazing was associated with increasing cover. This result is not in concordance with a study with manipulated sheep grazing after a prescribed fire (5.7 sheep ha^−1^ in four late summer and autumn months), where it was observed that the cover of dwarf shrubs was negatively affected by grazing, and the cover of sedges and grasses was positively affected (Damgaard et al. 2013). However, it is unfortunate that the Danish ecological surveillance program does not collect information on stocking rates or seasonality of grazing management, but only on whether grazing has taken place or not. This lack of resolution prevents a detailed analysis of the effect of the currently applied grazing on Danish wet heathland vegetation. It has been hypothesized that both grazing abandonment, which eventually may lead to a succession into a more woody vegetation, and overgrazing, where grass species are generally favored, may disfavor dwarf shrub species in wet heathlands (Hampton 2008). However, there is still considerable uncertainty as to the effect of grazing on wet heathland vegetation, and it will be beneficial if future grazing management actions monitor the effects and regulate stocking rates and seasonality using adaptive management procedures.

### Abiotic variables and soil pH

There were no significant effects of nitrogen deposition on soil pH (*γ*_*N*_, Table S2), whereas soil pH was found to be significantly higher on sandy soils compared to less sandy soils (*γ*_*S*_, Table S2). Based on these results, it was concluded that the effect of nitrogen deposition on plant community dynamics mainly was through direct effects, whereas the effect of soil type on plant community dynamics was both directly and indirectly mediated by the effect of soil type on soil pH. The observed insignificant effect of nitrogen on soil pH is in concordance with the earlier findings (Damgaard 2019), but the observed effect of soil type on soil pH is reversed in this study compared to the conclusions of the earlier study, which again suggests that relatively long time-series are needed for studying ecosystem dynamics. The soil acidification effects of nitrogen deposition are due either to nitrate leaching or removal of base cations from the system by nature management (Williams and Anderson 1999), and the present results could indicate that nitrate leaching or base cations removal from Danish wet heathlands is limited.

### Spatial variation

The short-scale spatial aggregation of the three species and the aggregate class of other higher plants were modelled by the parameter δ in the Dirichlet - multinomial mixture distribution (Appendix A). The estimated amount of short-scale spatial aggregation significantly increased the measurement uncertainty of the expected cover data compared to the case of randomly distributed plant species. If this over-dispersion of the pin-point cover data relative to the random expectation is not taken into account in the statistical model, then the signal to noise ratio will be severely upward biased and, most likely, will lead to erroneous conclusions (Damgaard 2013).

All species classes had significant regional geographic variation among the three assigned regions (Fig. 1). This result indicates that some of the large-scale spatial residual variation that could not be explained by the modelled environmental drivers may be explained by hitherto unexplored factors that differ among the three regions. In future studies, it will be important to understand the historical causes for the observed large-scale spatial variation in species abundance, e.g. by collecting and analyzing detailed accounts of site-specific nature management actions and hydrological data.

### Uncertainties and application of the model

The statistical modelling uncertainty was partitioned into measurement uncertainty and uncertainties due to the modelled spatial – and temporal processes. The most important source of measurement uncertainty was the plant cover measurement due to the significant small-scale spatial aggregation of plant species (see above), but the measurement uncertainty of nitrogen deposition, soil pH, soil C – N ratio, and soil type was also estimated and, thus, accounted for in the model. Generally, the structural uncertainty of the temporal processes was relatively small, whereas the large-scale spatial processes entail a higher degree of structural uncertainty.

One of the advantages of partitioning the different types of uncertainties in the SEM is the use of the fitted SEM for predictive purposes (Damgaard 2022), and since the fit of the temporal model was excellent, it is here suggested that the modeled environmental variables are sufficient for predicting the *average* plant community dynamics of wet heathlands. This optimistic conclusion is somewhat surprising in the light of several missing potentially important regulating variables, e.g. a direct measure of hydrology and previous natural management actions, which *a priori* were expected to be important factors in regulating plant community dynamics. However, caution and humbleness are required if the fitted model is used for generating local ecological predictions as input to a process of generating adaptive management plans for specific wet heathlands (Fig. 5), since the modelled environmental variables in this study may be correlated to unknown causal factors of plant community dynamics, contingent event with large effects, or causal factors where we do not have access to relevant environmental data. On the other hand, the modelling results provide important information to site managers on the relative importance of the different environmental factors. For example, since nitrogen deposition is invisible it is difficult to assess its effects without a statistical model and generally the effects of nitrogen deposition tend to be downplayed by site managers.

**Fig. 5.**
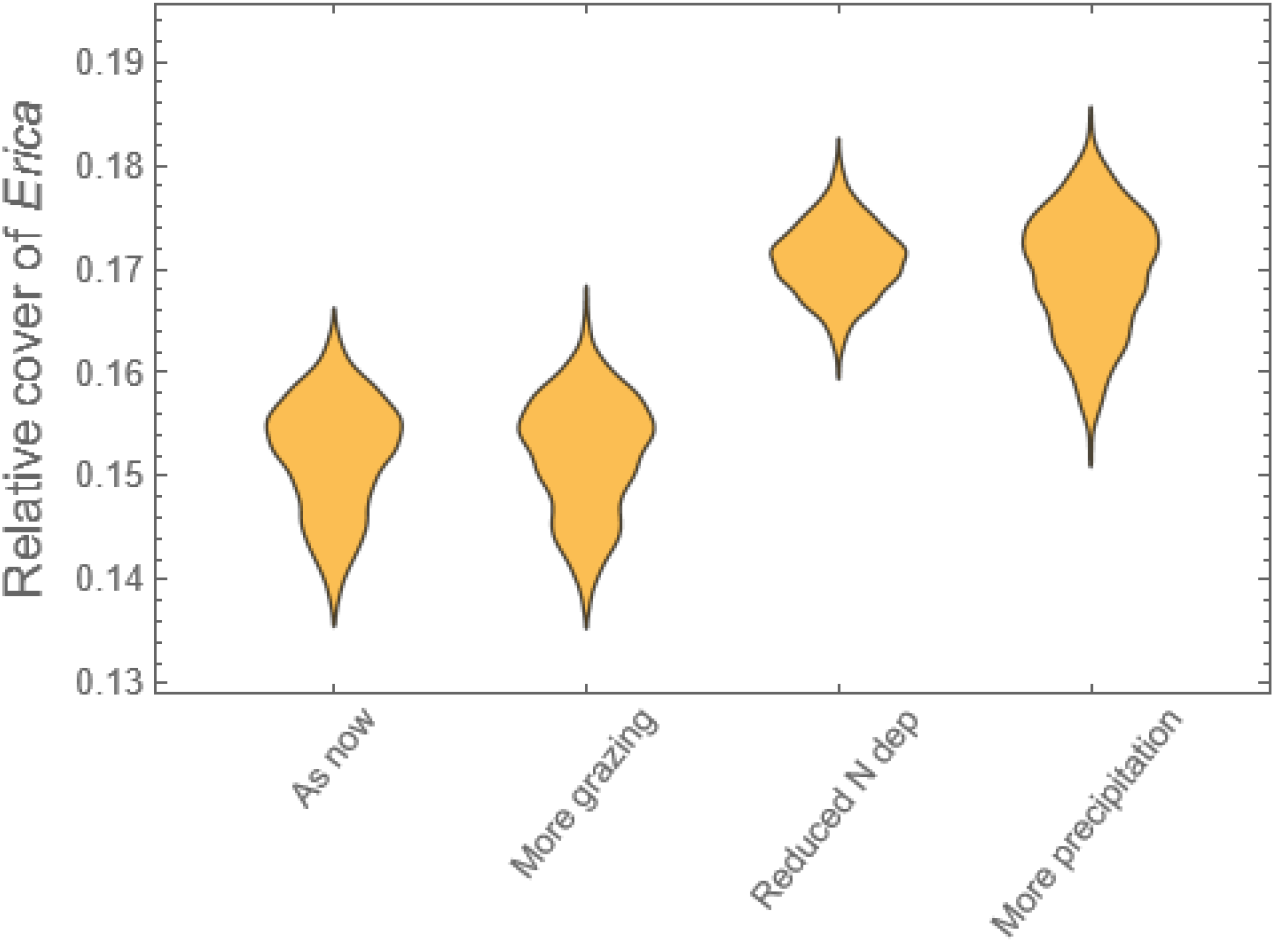
Predicted distribution of the cover of *Erica tetralix* for a specific wet heathland site (Musbæk Plantage in West Jutland) after five years under four different scenarios (Damgaard 2021). The initial cover of *E. tetralix* was 0.075 and scenarios were 1: As now, 2: more grazing, from 0.04 to 0.5, 3: reduced N deposition, from 14.1 kg N / ha year to 5kg N / ha year (note that it will require intensive mangement actions to reduce plant available N in the soil to the level expected at equlibrium under the reduced N deposition scenario), 4: more precipitation, from 881 mm / year to 1000 mm / year. The other environmental variables were pH in soil: 3.33, C-N ratio in soil: 25.7, soil texture (jb nr.): 1.1 (coarse sandy soil).

Moreover, the results suggest that the ratio between the two species *E. tetralix* and *M. caerulea* may be used as an indicator for the conservation status of wet heathlands.

## Electronic supplements

*The following electronic supplements may be downloaded from https://osf.io/SA8CF/*

Table S1 – Cover and change in cover of single species at wet heathlands

Table S2 - Marginal distribution of parameters

Table S3 – Proportion of variance explained

Fig. S1 – Pairwise scatter plot of variables

Fig. S2 - Dunn–Smyth residuals

Appendix A - Distribution of pin-point plant cover data

Appendix B - Spatio-temporal modelling

Appendix C - Estimating species cover and change in species cover

Appendix D – Mathematica notebook (a free reader may be downloaded from https://www.wolfram.com/player/)

## References

Aerts R, Berendse F. 1988. The effect of increased nutrient availability on vegetation dynamics in wet heathlands. Vegetatio. 76:63–69.

Aerts R, Berendse F, Caluwe Hd, Schmitz M. 1990. Competition in heathland along an experimental gradient of nutrient availability. Oikos. 57:310–318.

Bannister P. 1966. Erica tetralix l. J Ecol. 54(3):795–813.

Bobbink R, Loran C, Tomassen H. 2022. Review and revision of empirical critical loads of nitrogen for europe. German Environment Agency.

Damgaard C. 2012. Trend analyses of hierarchical pin-point cover data. Ecology. 93:1269–1274.

Damgaard C. 2013. Hierarchical and spatially aggregated plant cover data. Ecol Inform. 18:35–39.

Damgaard C. 2019. Spatio-temporal structural equation modeling in a hierarchical bayesian framework: What controls wet heathland vegetation? Ecosystems. 22:152–164.

Damgaard C. 2020. Measurement uncertainty in ecological and environmental models. Trends Ecol Evol. 35:871–873.

Damgaard C. 2021. Adaptive management plans rooted in quantitative ecological predictions of ecosystem processes: Putting monitoring data to practical use. Environmental Conservation. 1–6.

Damgaard C. 2022. Processes and predictions in plant ecological models: Logic and causality. EcoEvoRxiv.

Damgaard C, Irvine KM. 2019. Using the beta distribution to analyze plant cover data. J Ecol. 107:2747–2759.

Damgaard C, Nielsen KE, Strandberg M. 2017. The effect of nitrogen deposition on the vegetation of wet heathlands. Plant Ecol. 218(4):373–383.

Damgaard C, Strandberg MT, Kristiansen SM, Nielsen KE, Bak JL. 2014. Is erica tetralix abundance on wet heathlands controlled by nitrogen deposition or soil acidification? Environmental Pollution. 184:1–8.

Damgaard C, Thomsen MP, Borchsenius F, Nielsen KE, Strandberg M. 2013. The effect of grazing on biodiversity in coastal dune heathlands. J Coast Conserv. 17(3):663–670.

Damgaard C, Weiner J. 2021. The need for alternative plant species interaction models. Journal of Plant Ecology.

Degn HJ. 2006. Lyng og græs på randbøl hede 2005 - de store linier. Naturstyrelsen.

DMI. 2014. Average annual precipitation in the period 2001 to 2010 with a spatial resolution of 10 km Copenhagen: Danmarks Meteorologiske Institut.

Fremtidens klima i danmark. 2017. København: Danmarks Meteorologiske Institut; [accessed]. https://www.dmi.dk/klima/fremtidens-klima/danmark/.

Ellenberg H. 1979. Zeigerwerte der gefässpflanzen mitteleuropas. Scripta Geobotanica. 9(2. Ed).

Ellermann T, Andersen HV, Bossi B, Christensen J, Løfstrøm P, Monies C, Grundahl L, Geels C. 2012. Atmosfærisk deposition 2011 – novana. Aarhus: Nationalt Center for Miljø og Energi.

Ellermann T, Nygaard J, Christensen JH, Løfstrøm P, Geels C, Nielsen IE, Poulsen MB, Monies C, Gyldenkærne S, Brandt J et al. 2018. Nitrogen deposition on danish nature. Atmosphere. 9(11).

EU. 1992. Council directive 92/43/eec of 21 may 1992 on the conservation of natural habitats and of wild fauna and flora. In: Commission E, editor.

EU. 2013. Interpretation manual of european union habitats. Bruxelles: European Commission, DG Environment, Nature and Biodiversity.

Gimingham C. 1978. Calluna and its associated species: Some aspects of co-existence in communities. Plant Ecol. 36(3):179–186.

Gimingham CH. 1960. Biological flora of the british isles. No. 74. Calluna vulgaris (l.) hull. J Ecol. 48:455–483.

Gimingham CH. 1988. A reappraisal of cyclical processes in calluna heath. Vegetatio. 77(1/3):61–64.

Gimingham CH, Chapman SB, Webb NR. 1979. European heathlands. In: Specht, r.L. (ed), ecosystems of the world 9a, pp. 365–386. Elsevier, ansterdam..

Gimingham CH, Hobbs RJ, Mallik AU. 1981. Community dynamics in relation to management of heathland vegetation in scotland. Vegetatio. 46(1):149–155.

Grace JB, Anderson TM, Olff H, Scheiner SM. 2010. On the specification of structural equation models for ecological systems. Ecological Monographs. 80:67–87.

Granger CWJ. 1969. Investigating causal relations by econometric models and cross-spectral methods. Econometrica. 37(3):424–438.

Grant SA, Hunter RF, Cross C. 1963. The effects of muirburning molinia-dominant communities. Journal of the British Grassland Society. 18:249–257.

Grant SA, Torvell L, Common TG, Sim EM, Small JL. 1996. Controlled grazing studies on molinia grassland: Effects of different seasonal patterns and levels of defoliation on molinia growth and responses of swards to controlled grazing by cattle. J Appl Ecol. 33:1267–1280.

Greve MH, Greve MB, Bøcher PK, Balstrøm T, Breuning-Madsen H, Krogh L. 2007. Generating a danish raster-based topsoil property map combining choropleth maps and point information. Danish Journal of Geography. 107:1–12.

Hampton M. 2008. Management of natura 2000 habitats. 4010 northern atlantic wet heaths with erica tetralix. Bruxelles: European Commission.

Levy EB, Madden EA. 1933. The point method of pasture analyses. New Zealand Journal of Agriculture. 46:267–279.

Lindquist B. 1931. Den skandinaviska bokskogens biologi. Svenska Skogsvårdsföeningens Tidskrift. 3:179–485.

Lykke IMØ, Strandberg M, Nielsen KE, Barfod A, Damgaard C. 2015. Strukturelle ligningsmodeller som beslutningsgrundlag indenfor naturvaltningen - et eksempel fra pleje af klokkelyng på våde heder. Silkeborg: DCE.

Løvschal M. 2021. Anthropogenic heathlands: Disturbance ecologies and the social organisation of past super-resilient landscapes. Antiquity Project Gallery.

Løvschal M, Damgaard CF. 2022. Mapping the ecological resilience of atlantic postglacial heathlands. J Appl Ecol. n/a(n/a).

Nielsen KE, Bak JL, Bruus M, Damgaard C, Ejrnæs R, Fredshavn JR, Nygaard B, Skov F, Strandberg B, Strandberg M. 2012. Naturdata.Dk - danish monitoring program of vegetation and chemical plant and soil data from non-forested terrestrial habitat types. Biodiversity & Ecology 4:375.

Kontrolovervågning af terrestriske habitatnaturtyper 2004 – 2022. Novana. 2024. Aarhus Universitet: DCE – Nationalt Center for Miljø og Energi; [accessed]. www.novana.au.dk.

Nygaard B, Damgaard C, Bladt J, Ejrnæs R. 2020. Fagligt grundlag for vurdering af bevaringsstatus for terrestriske naturtyper. Artikel 17-rapporteringen 2019. Aarhus Universitet, DCE – Nationalt Center for Miljø og Energi.

Ovaskainen O, Roy DB, Fox R, Anderson BJ. 2016. Uncovering hidden spatial structure in species communities with spatially explicit joint species distribution models. Methods in Ecology and Evolution. 7(4):428–436.

Pearl J. 2009. Causality. Models reasoning, and inferences. 2, editor. Cambridge Cambridge University Press.

Pärtel M. 2002. Local plant diversity patterns and evolutionary history at the regional scale. Ecology. 83:2361–2366.

Rutter AJ. 1955. The composition of wet-heath vegetation in relation to the water-table. J Ecol. 43:507–543.

Stace C. 1999. Field flora of the british isles. Cambridge Cambridge University Press.

Strandberg M, Damgaard C, Degn HJ, Bak JL, Nielsen KE. 2012. Evidence for acidification-driven ecosystem collapse of danish wet heathland. Ambio. 41:393–401.

Strandberg M, Nielsen KE, Damgaard C. 2018. Habitat monitoring reveals decreasing morlayer c:N ratios in danish heathlands. Ecological Indicators. 89:538–542.

Svenning J-C, Sandel B. 2013. Disequilibrium vegetation dynamics under future climate change. Am J Bot. 100:1–21.

Taylor K, Rowland AP, Jones HE. 2001. Molinia caerulea (l.) moench. J Ecol. 89(1):126–144.

Usher MB, Thompson DBA. 1993. Variation in the upland heathlands of great britain: Conservation importance. Biol Conserv. 66(1):69–81.

Watt AS. 1947. Pattern and process in the plant community. J Ecol. 35:1–22.

Williams BL, Anderson HA. 1999. The role of plant and soil processes in determining the fate of atmospheric nitrogen. In: Langan SJ, editor. The impact of nitrogen deposition on natural and semi-natural ecosystems. Dordrecht: Kluver

